# MMseqs2 desktop and local web server app for fast, interactive sequence searches

**DOI:** 10.1101/419895

**Authors:** Milot Mirdita, Martin Steinegger, Johannes Söding

## Abstract

The MMseqs2 desktop and web server app facilitates interactive sequence searches through custom protein sequence and profile databases on personal workstations. By eliminating MMseqs2’s runtime overhead, we reduced response times to a few seconds at sensitivities close to BLAST.

**Availability and implementation:** The app is easy to install for non-experts. Source code, prebuilt desktop app packages for Windows, macOS and Linux, Docker images for the web server application, and a demo web server are available at https://search.mmseqs.com.

## I. INTRODUCTION

The most popular sequence similarity search tool, BLAST [1, 2], has garnered ∼7000 citations per year during the last 5 years, attesting to the unremitting importance of sequence searches for biology. This popularity may be largely owed to the excellent web services with short response times despite fast-growing databases provided by the NCBI/NIH, which requires a huge compute infrastructure. The distributed approach of running searches *locally* on personal computers or IT platforms of companies and research groups allows for custom databases, high availability, and protects sensitive data. But web server applications for local homology searches are slow as they mostly rely on BLAST (e.g. [4, 8]). Here, we present an application software to search with protein and nucleotide sequences through custom protein sequence and profile databases using MMseqs2 [9], achieving response times of seconds instead of minutes at a similar sensitivity as BLAST.

## II. METHODS

### Reduced runtime overhead

MMseqs2 owes its sensitivity and speed mainly to its prefiltering stage, which rejects ∼99.99% of sequences. The prefilter uses a reverse *k*-mer index table for the target database and also requires matrices with similarity scores between 2-mers and between 3-mers to generate the lists of similar 7-mers [9]. Reading in the index table and computing these matrices on-the-fly takes ∼0.5 min of runtime overhead for each search. We reduced this to 0.05 s by (1) writing the index table, the matrices, and other precomputable data into a file if it does not yet exist, (2) mapping the file into main memory and using the operating system’s cache optimization to keep it in memory, and (3) optimizing I/O operations.

### Optimized sequence-to-profile search mode

The index table for profile databases stores, for each position in a profile, all *k*-mers with a profile similarity score above a threshold set by -s. The number of similar *k*-mers grows exponentially with *k*. To save memory, we chose a short *k* = 5 as default for this mode. We also added MMseqs2 utilities for creating profiles from multiple sequence alignments (MSAs) and converting between profile formats.

### Desktop and web server app

The new software consists of frontend, backend and workers executing MMseqs2. The frontend is built with web technologies, which allow it to be accessed through a web browser and also to be packaged as desktop GUI application with the Electron framework (electronjs.org). The backend provides a RESTful API and worker scheduling. In the desktop app, all components run locally on the user’s workstation. For web server use, the components are deployed as Docker containers. The app takes protein sequence databases in FASTA format and MSA databases in STOCKHOLM format and automatically precomputes the index tables. If nucleotide sequences are used as queries, MMseqs2 automatically predicts open reading frames and searches with the translated sequences. Search results are shown with a customized feature-viewer (github.com/caliphosib/feature-viewer) (**Figure 1A**) and can be downloaded in tabular BLAST format.

**FIG. 1.**
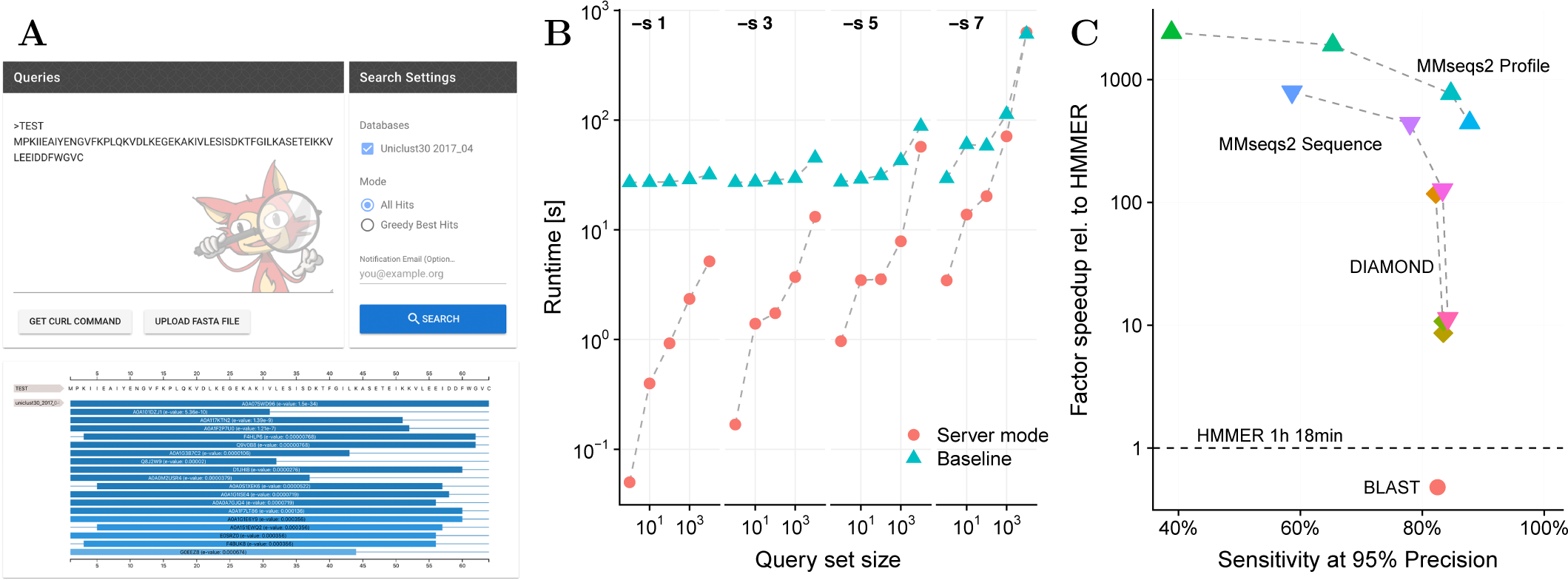
(**A**) Screenshots of the search interface and result visualization. (**B**) Runtime of searches with the baseline MMseqs2 (teal) and the new server mode (red) at four sensitivity settings (-s). (**C**) Domain annotation: Speedup versus sensitivity at 95% precision for MMseqs2 (at four sensitivity settings, -s 1, 3, 5, 7), DIAMOND (*default*, --sensitive, --more-sensitive) and BLAST. HMMER3 matches to Pfam domains are used as ground truth. The speed-ups exclude the times to format the databases.

## III. RESULTS

**Figure 1B** demonstrates the reduction of runtime overhead by comparing the runtimes of the MMseqs2 version without (“baseline”) to the new version with precomputations and memory mapping (“server mode”). Runtimes refer to searches with query sets of 1, 10, 100, 1000, and 10 000 sequences of average length 350 (sampled from the Uniclust30 database) through the Uniclust30 2017 10 database [7] with 13.5 million sequences, measured on a server with 2 Intel Xeon E5-2680 v4 CPUs with 14 cores each. The index table and matrix precomputation (∼3 min 40 s) is not included in the runtimes.

To test on metagenomics data the quality and speed of annotating Pfam domains, we built a test set by sampling 100 000 full-length sequences longer than 150 residues from our Marine Eukaryotic Reference Catalogue [10], clustering this set to 30% maximum pairwise sequence identity with MMseqs2, and sampling 10 000 sequences from the redundancy-reduced set. We annotated these sequences with PfamA 31.0 domains [6] using HMMER3 [5]. We then compared how well the sequence-sequence searches of MMseqs2, BLAST, and [3] DIAMOND and the sequence-to-profile searches of MMseqs2 could find the correct domain annotations. For the sequence-sequence search methods, we built a database from all sequences in PfamA.full MSAs and reported as E-value of a Pfam domain the E-value for the best-matching sequence from its MSA. We defined a search as true positive (TP) if the top match was annotated by HMMER3 with an E-value better than 10^*-*3^ and as false positive (FP) if the top match was not annotated with an HMMER3 E-value below 1. All other searches were considered ambiguous and ignored. For each method, we determined the E-value at which the precision TP/(TP+FP) is 95% and measured the sensitivity at that E-value.

As **Figure 1C** shows, MMseqs2 sequence-to-profile searches are ∼30 times faster than sequence-sequence searches with DIAMOND, MMseqs2 and BLAST and ∼300 times faster than HMMER3. The sensitivities and precision relative to HMMER3 are only lower limits, as any wrong HMMER3 match is not only considered true but causes the other tools’ correct matches to be considered false. Despite this, MMseqs2 sequence-to-profile searches reach 87% relative sensitivity at 95% precision, making them an attractive alternative to HMMER3 when speed is critical.

## IV. CONCLUSION

The desktop and web server app for MMseqs2 performs fast sequence searches at unprecedented speed-to-sensitivity trade-off on local computers. 1000 queries take only a minute to search through 15 million sequences of the Uniclust30 database, much faster than NCBI’s BLAST website. We hope the MMseqs2 app will empower also users unfamiliar with command line interfaces to perform fast and sensitive searches with their own sequence and profile databases.

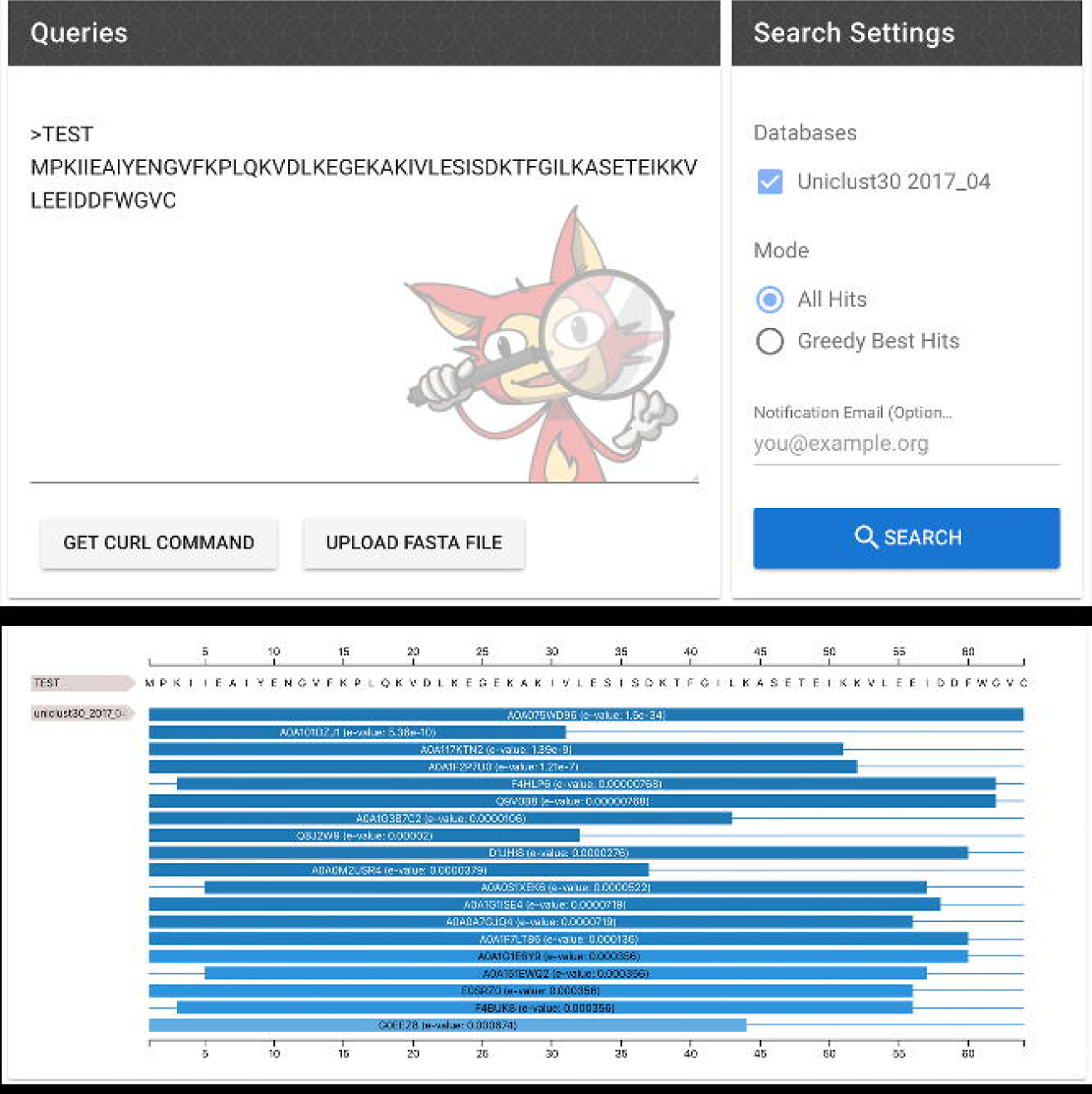

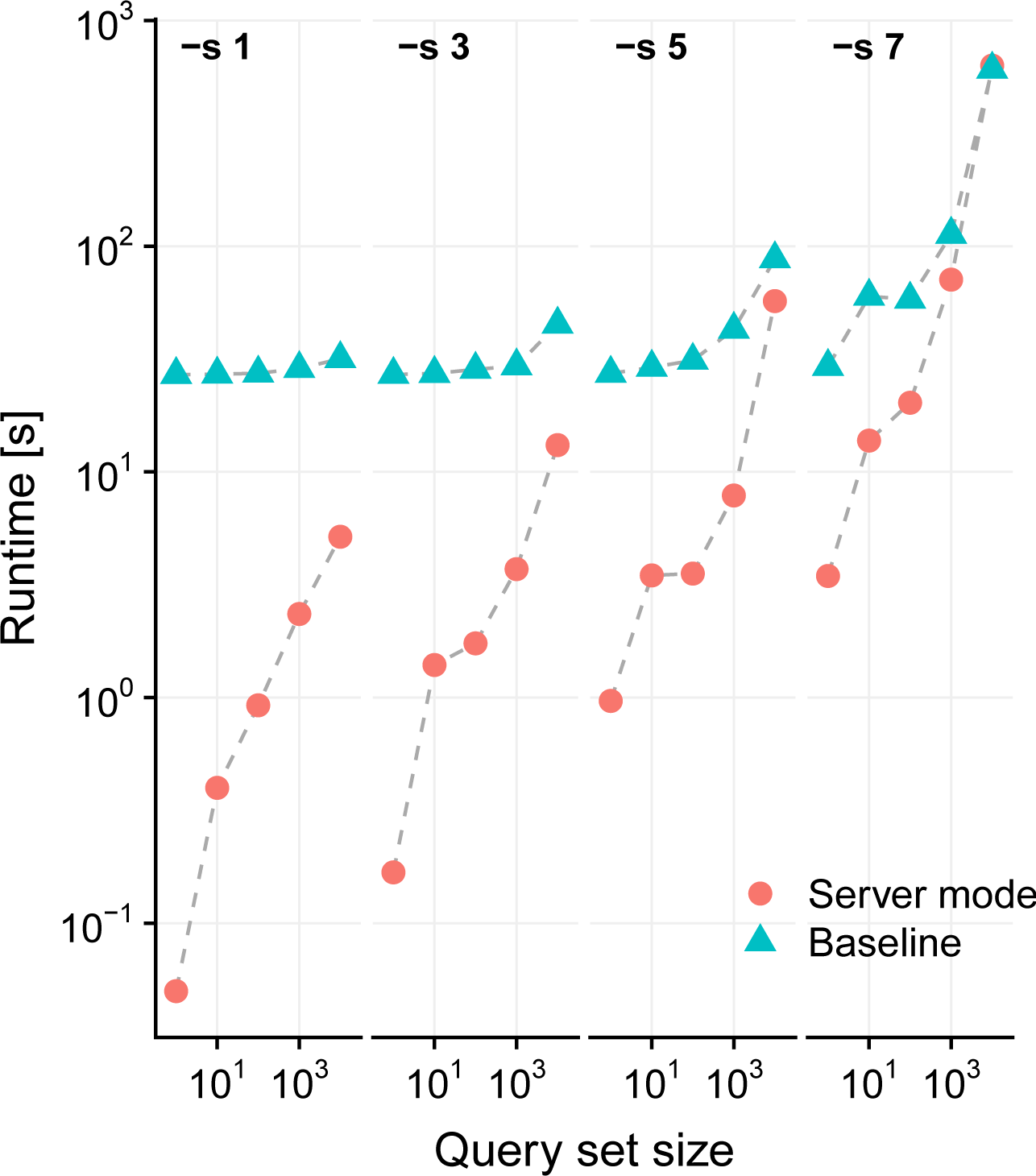

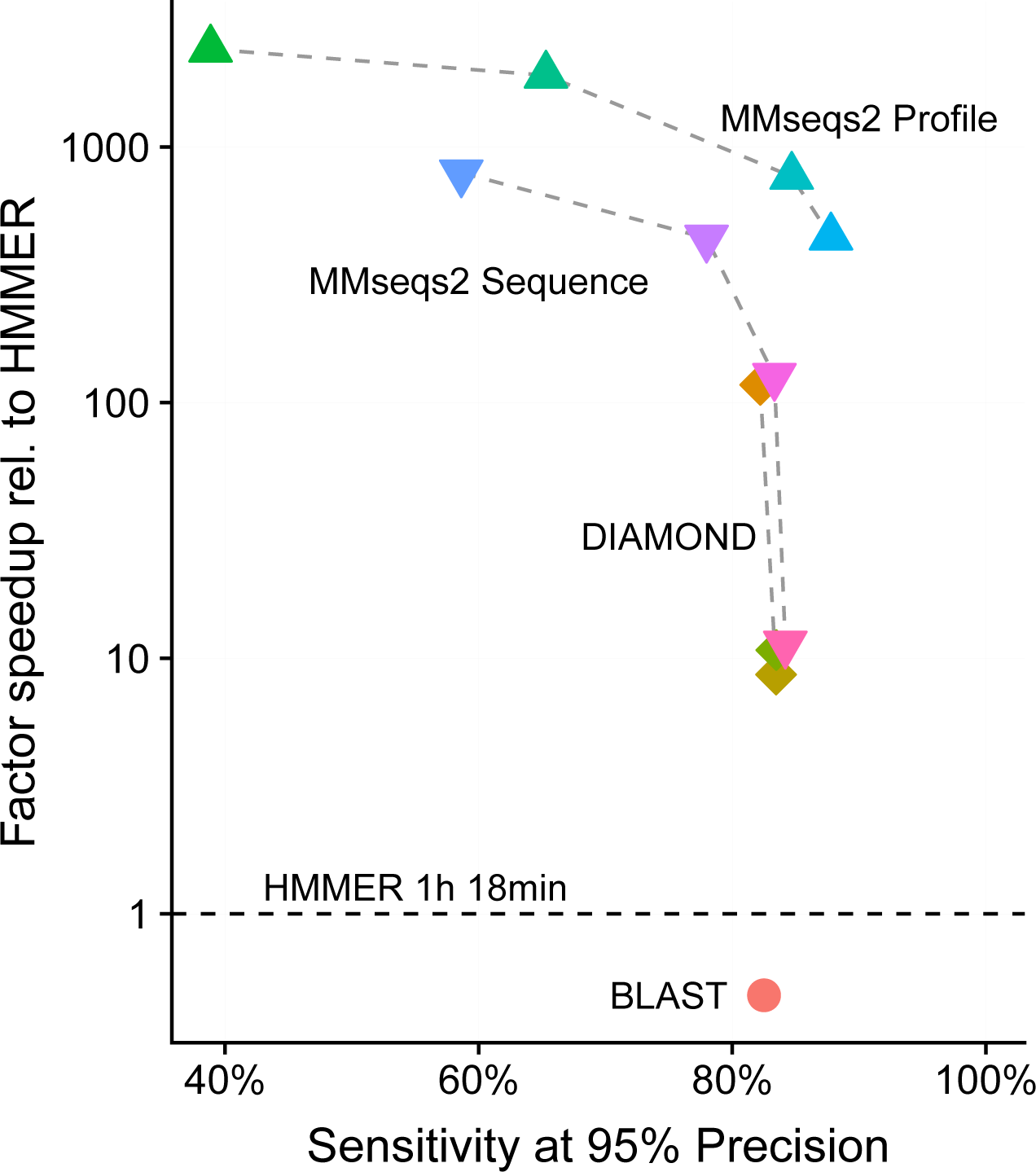

## ACKNOWLEDGEMENTS

We thank Yuna Kwon for crafting the “little Marv” mascot.

